# Paced Breathing With a Prolonged Inspiratory Period Increases Sympathetic Activity: A Heart and Brain Analysis

**DOI:** 10.1101/2024.02.23.581710

**Authors:** Miguel S. Joaquim, Carlos Moreira

## Abstract

Paced breathing exercises with prolonged exhalation have been commonly used to reduce stress and anxiety by stimulating parasympathetic activity. However, increasing sympathetic activity may also provide benefits such as increased alertness and energy levels. In this work, we investigate the physiological impact of an 80-second breathing exercise with a prolonged inspiratory period of 6 seconds followed by 2 seconds of exhalation on the sympathetic system. We collected raw two-channel prefrontal electroencephalography and photoplethysmography signals of 19 subjects using EMOTAI’s headband while performing the proposed exercise every workday for 2 weeks straight. Physiological metrics such as heart rate, heart rate variability, and absolute power of the brain waves were extracted before and during the exercise to measure its effectiveness. A marked increase in beta wave power was observed, along with a significant increase in both heart rate and heart rate variability. The cardiovascular results indicate that the proposed exercise effectively raised sympathetic activity. Simultaneously, the observed neural activity is consistent with that seen during focused attention and heightened mental processing.

## 1. INTRODUCTION

The voluntary modification of the breathing patterns is a powerful tool to modulate the Autonomic Nervous System (ANS), which regulates involuntary physiological processes such as heart rate, heart rate variability (HRV) and respiration. The ANS is composed of two complementary systems: the sympathetic nervous system, which prepares the body for stress, and the parasympathetic nervous system, which promotes relaxation (Betts et al., 2013; Shaffer et al., 2014).

Using paced breathing to help our body activate its “fight-or-flight” response (sympathetic activation) can be relevant in specific disease contexts (Kox et al., 2014; Thorp & Schlaich, 2015), as well as in healthy subjects during exercise, and physical and mental perturbations (Coote & Chauhan, 2016; Jordan & Tank, 2020; Sleight et al., 1995). More specifically, sympathetic activation in healthy individuals can have several benefits (Vijayalakshmi & Surendiran, 2005), including increased alertness (Pressman & Fry, 1989), enhanced cognitive performance, increased energy and motivation (Stancak et al., 1993), and improved immune functions.

In the present study, we propose an innovative breathing exercise with an increased inspiratory to expiratory ratio (i/e) expected to favour sympathetic activation in healthy individuals.

## 2. RELATION TO PRIOR WORK

Different paced breathing techniques were shown to modulate both the parasympathetic and sympathetic nervous systems (sympathovagal balance) (Bernardi et al., 2001; Zaccaro et al., 2018) with several studies successfully exploring the effectiveness of slow-breathing exercises in promoting parasympathetic stimulation (Bae et al., 2021; Komori, 2018; Laborde et al., 2022; Laborde, Hosang, et al., 2019; Laborde, Lentes, et al., 2019; Laborde et al., 2021; Magnon et al., 2021; You et al., 2021, 2022). However, it is still not yet completely understood to which extent the features of the paced breathing, other than respiratory rate, contribute to its effectiveness.

During inhalation, the vagal outflow is inhibited and favours sympathetic activation. In contrast, exhalation mediates vagal flow restoration due to parasympathetic activation. Therefore, controlling the relative timing of inspiration and expiration in breathing exercises (i/e ratio) seems to be an effective way to modulate the sympathovagal balance. A recent study (Laborde et al., 2021) showed that a deep and slow breathing pattern with a prolonged exhalation phase (low i/e ratio) stimulates the PNS for a higher period, which evidenced positive cognitive and physiological outcomes (Hoffmann et al., 2019; Laborde et al., 2022; Laborde, Hosang, et al., 2019; Laborde, Lentes, et al., 2019; Laborde et al., 2021).

The most common method for assessing the impact of breathing exercises on parasympathetic activity is through the respiratory sinus arrhythmia (RSA). RSA is responsible for the modulation of both heart rate and HRV in accordance with the breathing pattern (Strauss-Blasche et al., 2000). Thus, HRV measurements can be used as indirect indicators of parasympathetic activity (Junichiro Hayano, 2019). In another study (Van Diest et al., 2014), participants practising slow breathing with a longer exhalation time (low i/e ratio) showed increased HRV and stress reduction.

Altered breathing patterns have also been proven to impact cortical functions. Accordingly, studies have explored the use of electroencephalography (EEG) to assess the effects of paced breathing on the following brain waves: delta (*δ*, 0.5–4 Hz), theta (*θ*, 4–8 Hz), alpha (*α*, 8–13 Hz), beta (*β*, 13–30 Hz), and gamma (*γ, >* 30 Hz) (Teplan, 2002).

One of the first studies on paced breathing and EEG (Stancak et al., 1993) showed that EEG signals varied with breathing frequency, and that slower breathing was associated with decreased cortical arousal and increased relaxation. A similar study (Bušek & Kemlink, 2005) further established the relationship between breathing frequency and brain oscillations, with a decrease in the breathing frequency leading to increased total power in the anterior region. A randomized controlled trial (Cheng et al., 2018) found that deep breathing induced a significantly larger frontal theta power while the beta power decreased. The majority of studies on slow-paced breathing have consistently reported increases in theta and alpha bands, although this pattern was not observed in every case (Cahn & Polich, 2006; Zaccaro et al., 2018).

This study deviates from the typical approach of previous literature, which generally focuses on reducing sympathetic activity. Instead, we seek to examine the effectiveness of a breathing exercise with a prolonged inspiratory period whose objective is to increase sympathetic activity. With that purpose, we will be utilizing both photoplethysmography (PPG) and EEG metrics to monitor changes in heart rate, heart rate variability, and brainwave activity, respectively.

## 3. METHODOLOGY

### 3.1. Stimulus

In order to evaluate the physiological effects of prolonged inspiratory periods, a breathing exercise was selected with a 6-second inhalation followed by a 2-second exhalation. The breathing frequency was 7.5 cycles per minute (cpm), and the i/e ratio was 3/1. The exercise lasted for a total of 80 seconds, inspired by similar exercises available in mobile applications (Breathly, 2023; Breathwrk, 2023; Oak, 2023).

The proposed breathing exercise was implemented using the EMOTAI App (EMOTAI, 2023). The app included both visual and auditory pacers to guide the subjects, while also providing real-time biofeedback by plotting the subject’s heart rate alongside the visual pacer. Subjects were instructed to follow the pacer with their heart rate as seen in figure 1.

**Fig. 1:**
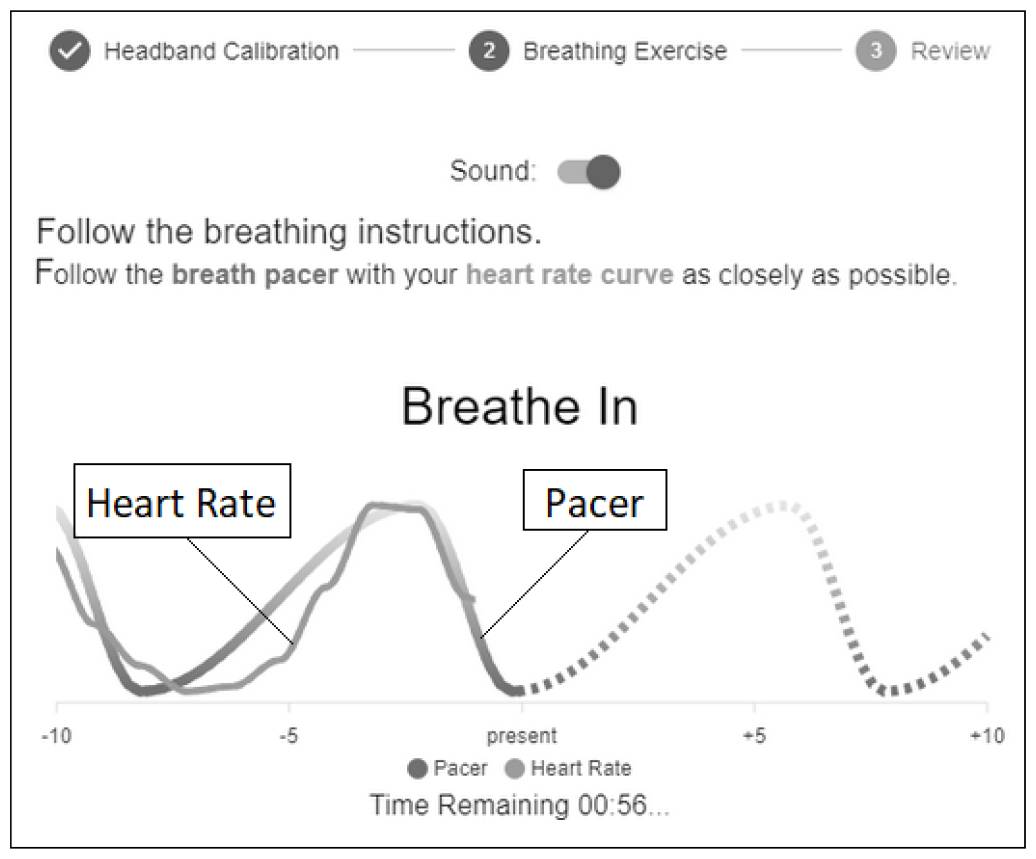
Screenshot of an ongoing breathing exercise in the EMOTAI App.

### 3.2. Data acquisition

In this study, raw EEG and raw PPG data from 19 subjects (12 men, 7 women, mean age = 37.71, age range = 24-56) were collected with a sampling rate of 100 Hz using EMOTAI’s wearable headband which is built upon the BITalino device (Batista et al., 2019). The wearable contains an optical PPG ear-clip sensor and 2 EEG prefrontal channels (Fp1 and Fp2).

Each participant was given a headband and access to EMOTAI’s software, which allowed them to do the exercise. Participants were shown how to properly place the sensors and were introduced to the breathing exercise through an in-app tutorial. The study lasted for 2 weeks and subjects were to perform at least one breathing exercise per day during each workday (10 days total), preferably in the morning.

In order to ensure the quality of the data collected during the exercises, the software had a calibration menu which prevented the subjects from starting the exercise unless the signal quality of both the PPG and EEG sensors was good, non-noisy and stable. Additionally, at least 15 seconds of raw data with good signal quality had to be collected for every sensor. The portion of initial data with good signal quality was later used as a baseline for further analysis. The full protocol can be seen in figure 2.

**Fig. 2:**
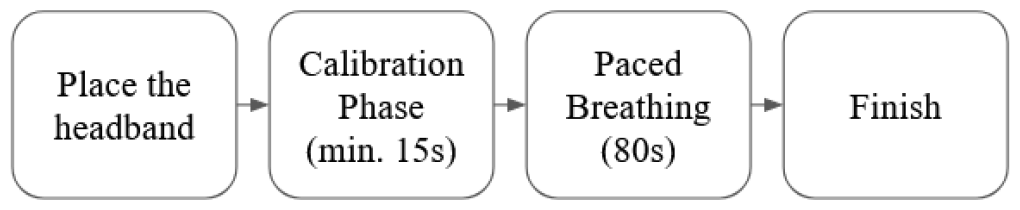
Flowchart of the protocol from placing the headband to the end of the exercise.

### 3.3. Proposed features

Aiming to understand if the proposed breathing exercise had any significant physiological effect on the subjects, several metrics were extracted from the raw data collected during both the calibration phase and the exercise itself.

#### 3.3.1. PPG metrics

From the data collected through the PPG sensor, both the heart rate and the heart rate variability were extracted. The heart rate values were computed every second using a 6-second window with a 5-second overlap The HRV values were computed using the RMSSD formula, with each value being calculated using 15-second windows (Munoz et al., 2015) and updated every 15 seconds with no overlap.

Overall baseline values for both heart rate and HRV were calculated using the median of the data from the calibration phase. This approach was also used to calculate an overall exercise value of the HRV, as the median is better at mitigating the impact of outliers than the mean. However, for the overall exercise value of the heart rate, a linear regression was performed using all the heart rate values measured throughout the exercise, and the considered aggregate value was the last one of the regression. This was done to compensate for the unique oscillatory pattern demonstrated by the heart rate due to the respiratory sinus arrhythmia.

#### 3.3.2. EEG metrics

The data collected from the EEG sensor was used to extract the frequency domain absolute powers of the bandwidth of interest, which include delta, theta, alpha, beta, and gamma. The power values were computed individually for each channel (Fp1 and Fp2) and each wave by applying the fast Fourier transform (FFT) algorithm to windows containing 128 samples (1.28 seconds) of raw EEG data (Ouyang et al., 2022). These windows were updated every second with a 28-sample overlap.

To improve the inter-subject comparability, the power of each brain wave for each EEG channel was normalized based on the sum of the power of all brain waves for that specific channel. The normalized brain waves from both EEG channels were combined into a single channel through averaging. Once again, overall baseline and exercise values were computed for all the normalized brain waves using the median.

### 3.4. Data Processing

When all subjects finished participating, a total of 147 recordings had been obtained. Every subject did at least 3 breathing exercises over the course of 10 workdays.

#### 3.4.1. Handling corrupted data

The first step in the data processing pipeline was checking the quality and integrity of the stored raw data by discarding recordings with a duration shorter than expected (80 seconds) or corrupted data. Only 2 recordings were discarded in this stage.

#### 3.4.2. Paced breathing detection

The next filtering criterion involved verifying if the subjects were accurately following the given instructions. This was achieved by exploiting the fact that, due to the RSA, the pattern of the heart rate should mimic the visual pacer when performing the exercise correctly (Strauss-Blasche et al., 2000). This behaviour can be seen in figure 3.

**Fig. 3:**
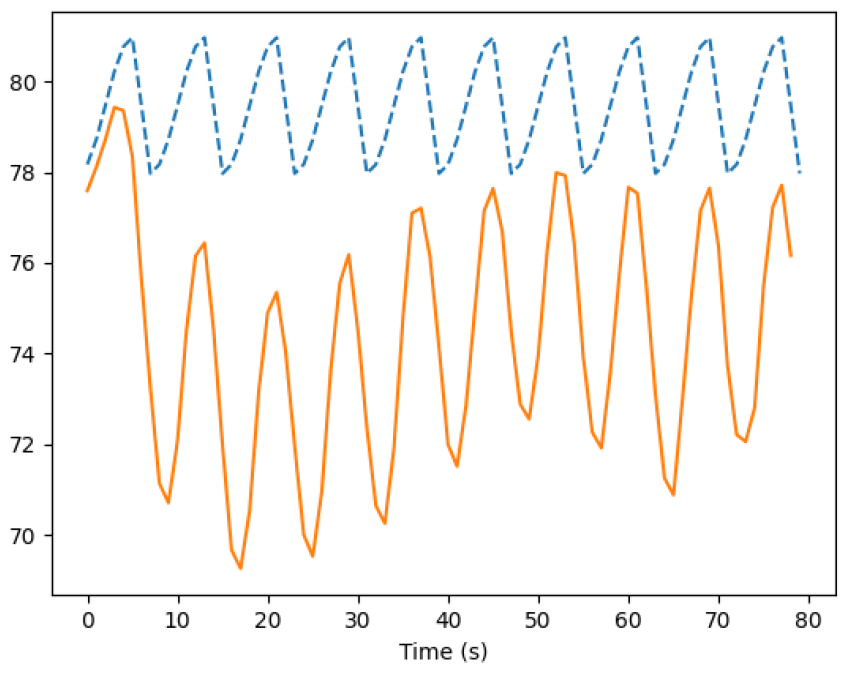
Average heart rate (—; beats per minute) across all analysed recordings and pacer (----; arbitrary units) during the breathing exercise.

Therefore, the following features were considered:

- The absolute value of the Pearson correlation coefficient (*r*) between the heart rate and the pacer for the full exercise. To account for any potential phase difference between them, a range of delay values between -3 and +3 seconds was considered with a 1-second step. A total of 7 correlation coefficients were calculated. The maximum absolute value of all these coefficients was used as the final correlation.
- The detected breathing frequency during the exercise. By analysing the median time difference between consecutive peaks in the heart rate plot caused by the RSA when transitioning from inhalation to exhalation, it was possible to estimate if the subjects were breathing at the target frequency of 7.5 cpm.

Regarding the Pearson correlation coefficient, the chosen threshold was |*r*| *≥* 0.20 as it was physically impossible to fully replicate the visual pacer and the subjects may not have had a consistent performance throughout the full exercise. Moreover, only recordings with an observed breathing frequency between 6 and 9 cpm (*M* = 7.40, *SD* = 0.36) were accepted. The goal was to only exclude recordings without any signs of the instructions having been followed properly.

After applying these criteria, 133 unique recordings were available for further analysis.

#### 3.4.3. Outlier Removal

Afterwards, the z-score (*z*) was calculated for the overall values of the proposed metrics obtained for both baseline and exercise. Outlier removal was done separately for PPG and EEG metrics to prevent one metric’s outliers from affecting the analysis of the other. Any recording containing any metric with |*z*| *≥* 3.0 was considered an outlier and discarded. After filtering, 123 unique recordings were available for PPG analysis, and 120 unique recordings were available for EEG analysis. This approach was crucial to ensure that the analysis of each sensor was independent and reliable, leading to more accurate results.

#### 3.4.4. Statistical Analysis

Finally, it was analysed if the breathing exercise had any statistically significant impact on the physiological parameters of the subjects. A comparison was made for all metrics between their exercises and respective baseline values using the Wilcoxon signed-rank (Woolson, 2007) (**n** = 123 for PPG metrics and **n** = 120 for EEG metrics) with *α* = 0.05. Additionally, the Holm-Bonferroni method (Aickin & Gensler, 1996) was used to minimize type I errors.

## 4. RESULTS

### 4.1. PPG analysis

#### 4.1.1. Heart Rate

Starting with the results for the heart rate, a statistically significant increase (*p<*.001, *T*=739) of the median heart rate values was observed. During the baseline, the median heart rate value across all recordings was 75.8 *±* 12.2 beats per minute (bpm) and increased to 80.3 *±* 13.1 bpm by the end of the exercise, corresponding to a 5.9% increase.

#### 4.1.2. Heart Rate Variability

When it comes to the HRV, a statistically significant increase (*p<*.001, *T*=997) of the median RMSSD was also registered. The median RMSSD during the baseline was 37.9 *±* 23.3 ms, increasing to 59.2 *±* 25.5 ms during the exercise, which corresponds to a 56.2% increment.

### 4.2. EEG analysis

All the results regarding the effects of the proposed breathing exercise on the bandwidth of interest of the EEG signal are summarized in table 1.

**Table 1:**
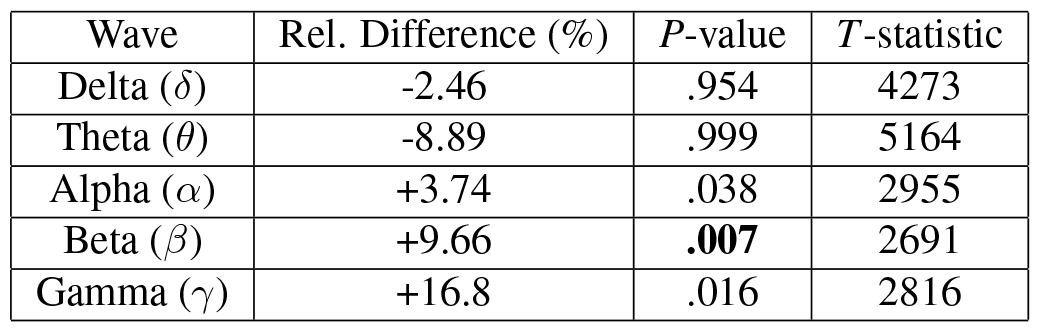
Percentage relative difference of the median relative power of the brain waves between exercise and baseline. Significant results are in **bold**.

It is possible to verify that the delta (*δ*), theta (*θ*) and alpha (*α*) waves exhibited no considerable difference before and during the breathing exercise. However, there was a statistically significant increase in the median values of the beta (*β*) waves during the exercise. Evidence also suggests that the gamma (*γ*) waves can potentially increase as an outcome.

## 5. DISCUSSION

Beginning with the analysis of the results from the PPG signal, a significant change in the HRV metric was detected. These findings are consistent with prior research studies (Laborde et al., 2021; Van Diest et al., 2014) that have employed paced breathing as a means of eliciting physiological responses in subjects.

In contrast with the commonly used breathing exercises with prolonged expiratory periods (low i/e ratio) that are used to induce relaxation, in this study a breathing exercise with a prolonged inspiratory period (high i/e ratio) was proposed. Still, despite this difference, it was possible to observe a noticeable impact of the exercise on the RSA, resulting in increased HRV values. A higher HRV, for instance, is associated with a better health status and a lower risk of diseases (Shaffer et al., 2014).

On the other hand, the results regarding heart rate differed from those observed in the exercises with slower paced breathing (Bernardi et al., 2001; Song & Lehrer, 2003). In our study, a significant heart rate increase was detected by the end of the breathing exercise. This pattern suggests an increased sympathetic activation which, in healthy amounts, correlates with improved alertness, energy levels and enhanced cognitive performance (Jordan & Tank, 2020; Vijayalakshmi & Surendiran, 2005).

When it comes to the results from the EEG, the relative power of delta (*δ*), theta (*θ*) and alpha (*α*) waves of the participants exhibited no significant difference during the exercise in comparison to the baseline measurements. This implies that the proposed exercise did not have any noticeable impact on these waves. As shown in previous studies, these low-frequency brain waves tend to increase in the prefrontal cortex during breathing exercises for anxiety reduction (Cahn & Polich, 2006). These patterns occur as the power of theta and alpha waves are positively correlated with states of drowsiness and relaxation (Marzbani et al., 2016).

However, a meaningful increase was observed in the relative power of the beta (*β*) waves with the gamma (*γ*) waves also showing that they can be potentially affected. The role of these brain waves in the prefrontal cortex has been shown to be associated with tasks requiring focused attention and higher mental effort, such as solving complex problems or performing creative tasks (Kropotov, 2010; Marzbani et al., 2016).

In the end, the registered relative power increase in the high-frequency brain waves coupled with the elevated heart rate and HRV levels during, and by the end of the exercise, demonstrate the existence of a link between breathing exercises with longer inspiratory periods (high i/e ratio) and an improved level of alertness or mental arousal among the subjects.

## 6. CONCLUSIONS

For a very long time, people have been using paced breathing exercises as a tool to induce various physiological effects. These exercises were the subject of numerous studies to scientifically validate their benefits and effects on well-being. While successful, previous literature has predominantly focused on the application of these exercises in reducing stress and anxiety by stimulating parasympathetic activity.

Based on the limited amount of studies targeting physiological effects other than relaxation, this study attempted to shed additional light on the topic by exploring a novel breathing technique aimed at increasing sympathetic activity to promote healthy levels of alertness and mental arousal.

In order to achieve that goal, participants performed a paced breathing exercise with a 6-second inhalation followed by a 2-second exhalation, for a total duration of 80 seconds, with at least 1 exercise per day for 10 weekdays straight. A significant increase in both heart rate and HRV was observed, together with a marked increase in the relative power of the beta (*β*) brain wave in the prefrontal cortex. The observed EEG patterns match the ones that occur during tasks that demand focused attention and mental processing, thus indicating that the proposed exercise successfully elicited the intended physiological effects.

Although existing studies support the benefit of paced breathing exercises, many questions remain unanswered. Further research is crucial to determine the optimal duration and the i/e ratio of these exercises. The number of subjects should also be as high as possible in order to ensure the generalizability of the results. Finally, it is necessary to investigate how long the effects of the exercise linger for after it has ended.

